# Neutron Scattering Analysis of *Cryptococcus neoformans* Polysaccharide Reveals Solution Rigidity and Repeating Fractal-like Structural Patterns

**DOI:** 10.1101/2023.09.22.559017

**Authors:** Ziwei Wang, Susana C. M. Teixeira, Camilla Strother, Anthony Bowen, Arturo Casadevall, Radamés JB Cordero

## Abstract

*Cryptococcus neoformans* is a fungal pathogen that can cause life-threatening brain infections in immunocompromised individuals. Unlike other fungal pathogens, it possesses a protective polysaccharide capsule that is crucial for its virulence. During infections, *Cryptococcus* cells release copious amounts of extracellular polysaccharides (exo-PS) that interfere with host immune responses. Both exo-PS and capsular-PS play pivotal roles in *Cryptococcus* infections and serve as essential targets for disease diagnosis and vaccine development strategies. However, understanding their structure is complicated by their polydispersity, complexity, sensitivity to sample isolation and processing, and scarcity of methods capable of isolating and analyzing them while preserving their native structure. In this study, we employ small-angle neutron scattering (SANS) and ultra-small angle neutron scattering (USANS) for the first time to investigate both fungal cell suspensions and extracellular polysaccharides in solution. Our data suggests that exo- PS in solution exhibits collapsed chain-like behavior and demonstrates mass fractal properties that indicate a relatively condensed pore structure in aqueous environments. This observation is also supported by scanning electron microscopy (SEM). The local structure of the polysaccharide is characterized as a rigid rod, with a length-scale corresponding to 3 to 4 repeating units. This research not only unveils insights into exo-PS and capsular-PS structures but also demonstrates the potential of USANS for studying changes in cell dimensions and the promise of contrast variation in future neutron scattering studies.

## INTRODUCTION

*Cryptococcal* meningitis is a deadly fungal infection caused by *Cryptococcus neoformans*, one of the leading causes of death in HIV/AIDS patients in sub-Saharan Africa^1,2^. Within brain tissues, the fungus secretes copious amounts of polysaccharide (exo-PS) to the cerebrospinal fluid, believed to cause elevated intracranial pressure and disruption of an effective immune response^2,3^. The fungal cell is encased by a thick capsule composed of polysaccharide (capsular-PS), which protects from the host’s immune defense mechanisms^4^. Both the exo-PS and capsular-PS are mainly comprised of glucuronoxylomannan (GXM), which is formed by an α-1,3-linked mannan backbone with β-1,2-linked glucuronic acid and β-1,2- or β-1,4-linked xylose as its branching residues that contribute to serological diversity. GXM molecules are assembled from six structural units (M1-6), referred to as triads, featuring a glucuronic acid residue every third mannose along with varying xylose substitutions (Figure 1). Due to its high water-content (over 95% of total mass and volume), the PS capsule is highly susceptible to the dehydration steps employed in high-resolution microscopy or lyophilization, which disturbs the native structure^5,6^.

**Figure 1.**
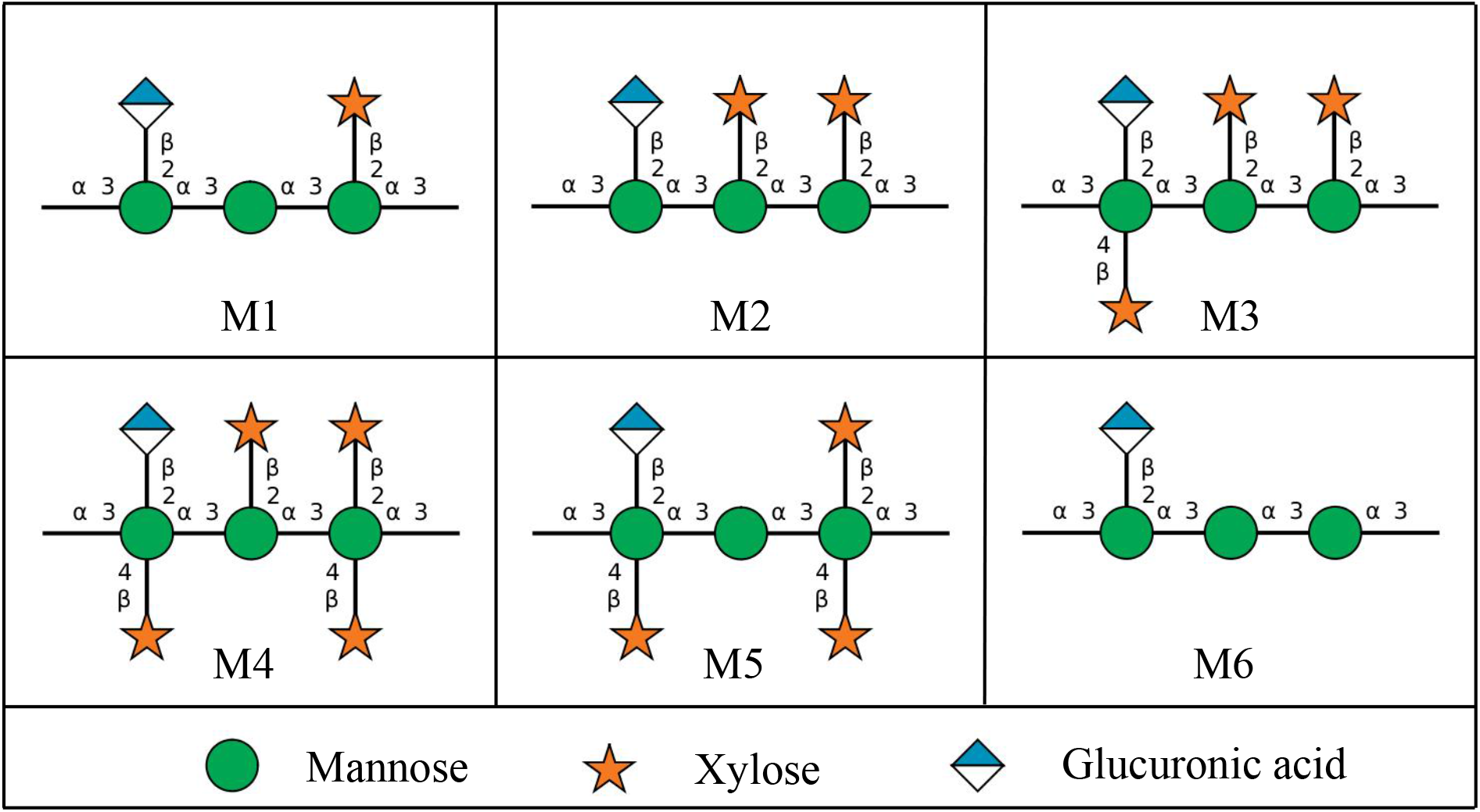
The six Glucuronoxylomannan (GXM) motifs that build up fungal exo-PS and capsular- PS. GXM is composed of a combination of six repeating units (M1-6), defined by a glucuronic acid (GlcA) residue every 3rd mannose with varying xylose substitutions. These 6 motifs of GXM in various combinations correlate to different serotype activities^19^. *C. neoformans* H99 serotype A has a dominant M2 motif in exo- and capsular-PS^20^. Polysaccharide molecules can be heteropolymers composed of more than one triad^17^. Image created with DrawGlycan-SNFG^21^.

*C. neoformans* exo-PS and capsular-PS are key virulence determinants and targets for the immune system, vaccine design, and monoclonal antibody (mAb) treatments. Diagnosis of *Cryptococcus* infection primarily relies on culturing the organism and antigen detection of shed PS^7,8^. Although both exo-PS and capsular-PS are predominantly composed of GXM, they exhibit distinct physicochemical properties and mAb reactivity^8^. Despite their significance in disease and diagnosis and the known chemical composition of the GXM, the macro- and supramolecular assembly of the *Cryptococcal* PS and the corresponding effects on epitope binding by antibodies remain largely unknown. Studying the correlation between nanoscale structure and macroscopic properties of the PS, and how both natural and experimental environments can trigger different assembly characteristics, is crucial for the design of diagnostic assays and strategies for vaccine development.

The large and complex PS heteropolymers display physicochemical characteristics that vary based on nutrient availability, chemical and physical environment, and cell age^8,9^. Consequently, experimental biases can be introduced by sample preparation protocols and the inherent limitations of measurement techniques. Previous research has primarily focused on PS isolated from culture supernatants using hexadecyltrimethylammonium bromide (CTAB) precipitation or filtration, and capsular-PS extracted via dimethylsulfoxide (DMSO) extraction and ionizing radiation-induced PS ablation, leading to nominally de-capsulated cells (residual capsular-PS may remain). A variety of techniques, such as static (SLS) and dynamic light scattering (DLS), zeta potential measurements, optical tweezers-based elastic modulus assessments, and solution viscosity analyses, have been employed to investigate exo-PS and capsular-PS structure^10,11^. SLS and DLS analyses suggest that the polysaccharide molecules are branched, a characteristic that influences immune reactivity and modulation^12–15^. Existing experimental data is consistent with capsular-PS polymers in molar mass ranges of 1-7 MDa, radii of gyration (*Rg*) ranging from 150-500 nm, hydrodynamic radii (*Rh*) ranging spanning 570-2000 nm, contingent on the specific experimental method used^10,13,16,17^. Encapsulated and nominally de-capsulated cells were previously investigated by powder X-ray diffraction, where broad peaks in the range of 1.46 Å^-1^-1.51 Å^-1^ of momentum transfer vector *q*, were attributed to a repeating structural motif arising from inter-molecular interactions mediated by divalent metals and glucuronic acid residues, and/or possibly gelated PS organization^18^. To achieve a detailed characterization of the relationship between the structures of exo-PS and capsular-PS and their corresponding functions at different stages of infection, measurement capabilities that preserve PS conformation are necessary.

In this study, we used neutron scattering analyses, light microscopy, and scanning electron microscopy to probe the structure of exo-PS, intact fungal cells, and gamma-irradiated nominally de-capsulated cells in solution (**Figure 2**). Neutron scattering accesses a broad range of structural features without causing radiation damage, enabling data collection on the same samples across small-angle (SANS) and ultra-small angle (USANS) neutron scattering regimes at various temperatures and concentrations. This approach has been previously used for studies of polysaccharides, such as arabinoxylans by Yu *et al.*^22^, and minimizes sample discrepancies, while the use of varying percentages of D2O in buffers allows for contrast variation. Given the considerable size of the cells (micrometers in diameter), the USANS regime is crucial, whereas the analysis of intrachain structures is conducted within the SANS regime.

**Figure 2.**
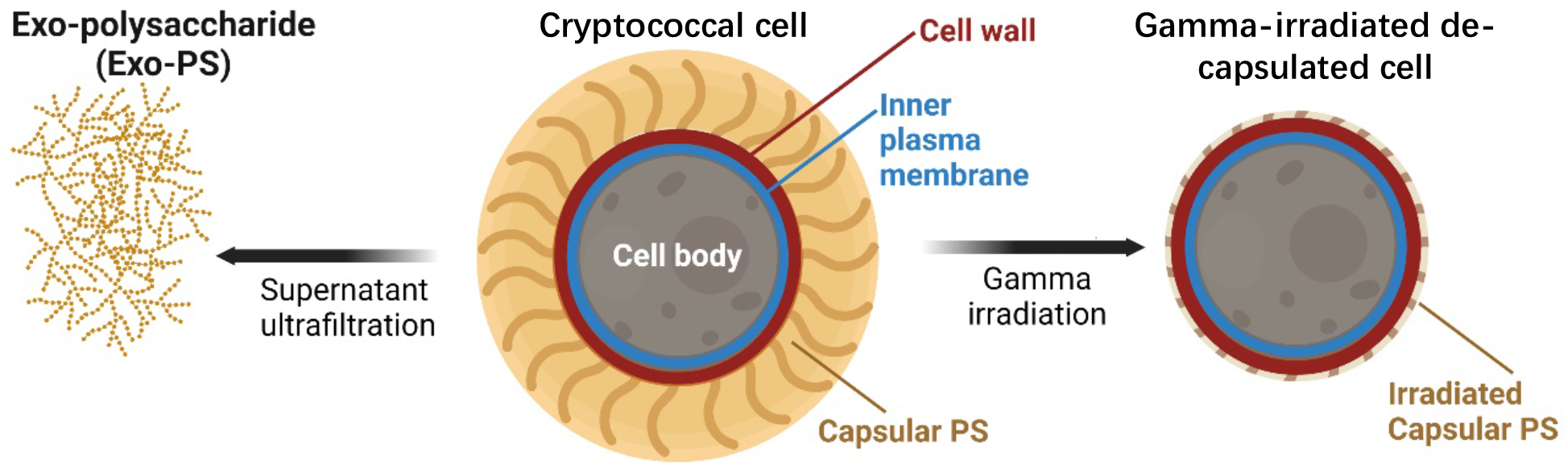
Schematic diagram of the samples investigated: exo-PS, *cryptococcal* cell, and gamma-irradiated de-capsulated cell. Exo-PS refers to the secreted polysaccharide, while capsular-PS refers to the highly hydrated polysaccharide surrounding the capsule of the intact cell. Gamma-irradiated capsular-PS refers to the remaining polysaccharide thin capsule surrounding the irradiated cells. Image created with BioRender.com.

## MATERIALS AND METHODS

### Fungal Growth

*Cryptococcus neoformans* Serotype A strain H99 was inoculated in 20 mL of Sabouraud dextrose broth and grown with agitation (120 rpm) for 2 days at 30 °C. The cells were pelleted by centrifuging for 10 minutes at 3000 rpm, and resuspended in minimal media (15 mM dextrose, 10 mM MgSO_4_, 29.3 mM KH_2_PO_4_, 13 mM glycine, and 3 μM thiamine-HCl, adjusted to pH 5.5 using KOH; where “M” represents the SI unit mol/L). The washing process was repeated three times, and the cells were finally resuspended in minimal media to a density of 1×10^6^ cells/mL. Subsequently, the cells were inoculated into a 1 L culture of minimal media and incubated at 30 °C for 7 days. The cells were harvested via centrifugation at 4,000 rpm for 20 minutes. The supernatant was filtered using a 0.22 µm Millipore.

Capsular-PS was removed by exposing the whole cells to 40 minutes of gamma irradiation, using the method demonstrated by Maxson *et al.*^23^. The ionizing radiation was demonstrated to be effective in removing the capsular-PS^24^.

### Exo-PS Isolation

The exo-PS was isolated from the cell-free supernatant as previously described^25^. The supernatant was sequentially filtered with an Amicon membrane filter (100 kDa nominal molar mass cutoff) and the flow-through was then filtered using a 10 kDa membrane filter. The exo-PS accumulated on the 10 kDa membrane surface as a clear gel and was collected and dialyzed extensively against MilliQ-grade H_2_O or D_2_O (Cambridge Isotope Labs, 99.9% D). The H_2_O solutions provide scattering profiles at an additional contrast, while the samples prepared in D_2_O are expected to minimize the incoherent neutron scattering background contribution to the measured intensity profiles. Following dialysis, the exo-PS concentration was determined using a phenol sulfuric colorimetry assay^26^. Sample solutions were centrifuged for 2 minutes at 10,000 rpm to remove any debris. This process resulted in 10mg/mL exo-PS samples with a hydrodynamic radius *Rh* of 550-600 nm and relatively low polydispersity, quantified as 0.355 (**Figure S1**) by DLS coupled with a 90Plus/BI-MAS Multi-Angle Particle Sizing analyzer (Brookhaven Instruments Corp., NY, USA), as described by Frases *et al.*^16^. The prepared exo-PS was also resuspended at 1 mg/mL and 5 mg/mL. Based on the equation derived by Vadillo *et al*.^27^ (*c\**≈ 1.46/[*]) and the intrinsic viscosity ([*]) of PS from strain H99 in minimal media determined by Cordero *et al*.^13^, the overlap concentration *c\** for exo-PS is estimated as 5 mg/mL^28^.

### SANS and USANS Data Collection and Reduction

Neutron scattering data were collected at the National Institute of Standards and Technology (NIST) Center for Neutron Research (NCNR; Gaithersburg, Maryland USA). All samples were degassed for 10 minutes before data collection. SANS data were obtained from the 30-meter instruments NG7 and NGB, using a neutron wavelength *λ* of 6 Å and a wavelength spread *Δλ/λ* of 12.5 % for three sample-to-detector distances, to measure scattered intensities over a range of momentum transfer defined as:

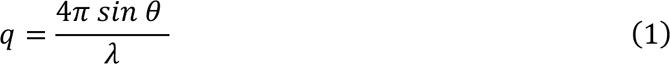

where *2θ* is the scattering angle measured. Focusing lenses were used for the longer wavelengths (8.4 Å on NGB, and 8.09 Å on NG7) to extend the lower *q* range to 0.001 Å^-1^ in the SANS regime^29^. Scattered neutrons were detected with a 64 cm × 64 cm 2D position-sensitive detector with 128 pixels × 128 pixels at a resolution of 0.508 cm/pixel. SANS data measured for solutions in H_2_O at concentrations of exo-PS up to 10 mg/mL show a flat intensity profile in the *q* range measured, indicative of insufficient contrast to provide a measurable signal above the strong incoherent scattering background from the hydrogen atoms in the buffer (**Figure S2**). SANS measurements were carried out on 1 mg/mL, 5 mg/mL, and 10 mg/mL of exo-PS solutions in D_2_O at three temperatures (22 °C, 30 °C, and 37 °C), controlled by a Peltier-driven sample changer, with 30 minutes of pre-equilibration at the desired temperature before data collection. The temperatures chosen include typical ambient experimental environments (22 °C), as well as the *C. neoformans* optimal growth (30 °C), and physiological temperatures (37 °C). No significant differences were observed between the SANS profiles of exo-PS solutions at the three measured temperatures (data not shown): the profiles overlapped well, within experimental error. A temperature of 30 °C was therefore chosen for data collection on the whole cells and gamma-irradiated cells in the SANS and USANS regime, to maintain consistency with the growth temperature of the whole fungal cells.

Slit-smeared USANS data were collected at the double-crystal diffractometer (Bonse-Hart) BT5 at the NCNR (*λ* = 2.4 Å, *Δλ/λ* = 6%)^30^, to cover a *q* range of 0.00003 Å^-1^ to 0.003 Å^-1^. USANS measurements were carried out on 10 mg/mL exo-PS in D_2_O at 30 °C to probe the presence of aggregates or finite size clusters or aggregates in the micrometer to hundreds of nanometers size range. USANS data were also collected on D_2_O and H_2_O solutions of the whole fungal cells, and gamma irradiated cells at the concentration of 1 × 10^!^cells/mL.

SANS and USANS data were reduced using the macro-routines developed for IGOR Pro at the NCNR^31^. Raw counts were normalized to a common neutron monitor count and corrected for empty cell counts, ambient background counts, and nonuniform detector response. The data obtained from the samples were placed on an absolute scale by normalizing the scattered intensity to the incident beam flux. Buffer-only reduced data were subtracted from SANS data on samples containing exo-PS or capsular-PS.

### Neutron Scattering Data Fitting

The SANS data for the exo-PS solutions in D_2_O were fitted using a modified empirical correlation length function that calculates scattering intensities as:

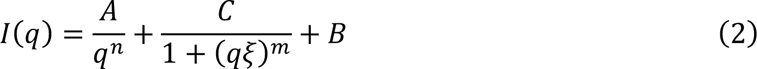

where the first term describes Porod scattering from pore clusters (exponent *n*) and the second term is a Lorentzian function describing scattering from the PS polymer chains (exponent *m*)^32^. The second term characterizes the PS/solvent interactions, and the two multiplicative factors *A* and *C* are, respectively, the Porod scale and the Lorentz scale. *ξ* is a correlation length for the PS chains, and *B* is a *q*-independent incoherent neutron scattering intensity background contribution to I(*q*). The calculated intensities from the correlation length model were smeared to match the instrumental pinhole smearing read from the reduced experimental data file. The exo-PS volume fraction and the *B* parameter were kept fixed throughout the fits. The fitting parameters and relevant information on the goodness-of-fit are available in the supporting information for the interested reader (**Table S1**). It was assumed that the entanglement of overlapping PS chains did not contribute significantly towards the SANS profiles.

### Light Microscopy

To measure the cell and capsule dimensions, whole cells and gamma-irradiated cells in D O and H O at 1 × 10^!^ cells/mL were imaged with an Olympus AX70 microscope, using the QCapture Suite V2.46 software for Windows. *Cryptococcal* cells were suspended in India Ink, which is excluded by the PS so that the capsule region will appear to be bright/empty. Cell dimensions were measured with ImageJ in pixels, and then converted to µm (152 pixels correspond to 50 µm for 40× magnification objective with 2×2 binning). Statistical analyses were performed using GraphPad Prism version 9.5.1 for Mac OS X, GraphPad Software, Boston, Massachusetts USA. Unpaired statistical analyses t-tests were done for the cell diameters and capsule thickness comparisons; the corresponding significance was stratified based on the probability that the results occur by chance, quantified as a probability through a percentage p- value, where 5% is equivalent to p = 0.05.

### Scanning Electron Microscopy

SEM of encapsulated *C. neoformans* yeast cells was done as previously described^13^. Briefly, the cells were fixed using a solution containing glutaraldehyde, sodium Cacodylate, sucrose, and MgCl_2_. After dehydration with ethanol, critical point drying was performed using liquid carbon dioxide. The dried samples were then sputter-coated with gold- palladium for improved conductivity. Finally, the prepared samples were visualized using a JEOL JSM6400 Scanning Electron Microscope at an accelerating voltage of 10 kV, enabling high- resolution imaging of the yeast cells. To analyze the fractal dimension of SEM images of whole cells and capsular-PS structures, we utilized the FracLac plugin of ImageJ (http://rsb.info.nih.gov/ij/plugins/fraclac/FLHelp/Introduction.htm). The FracLac algorithm quantifies the complexity of patterns in digital images, providing fractal dimensions data. The algorithm works by scanning the input micrographs using a shifting grid algorithm, which allows multiple scans from different locations on each image.

## RESULTS AND DISCUSSION

### SANS Analysis of Exo-PS Solutions

Based solely on the water-free composition, the neutron scattering length density (SLD) of polysaccharides in H_2_O is expected to range from 1×10^-6^ Å^-2^ to 2×10^-6^ Å^-2^ (the SLD of pure H_2_O is -0.56×10^-6^ Å^-2^)^33^. The flat scattering profiles observed in the exo- PS samples measured in H2O are consistent with a high hydration state and a significant contribution of incoherent scattering originating from the hydrogen atoms within the samples (**Figure S2**). In the case of exo-PS solutions in D2O, there is still a relatively strong incoherent neutron scattering background observed in the reduced SANS data due to the non-liable hydrogen atoms (**Figure 3**). The predominant M2 motif expected in our sample consists of three mannoses, two xyloses, and one glucuronic acid^20^, containing a significant number of labile hydrogens that can exchange against D2O during dialysis. However, non-labile and solvent-inaccessible hydrogen atoms that remain at varying concentrations are responsible for the differences in background intensities observed at high *q* in the SANS data for samples at 1 mg/mL, 5 mg/mL, and 10 mg/mL exo-PS (**Figure 3**).

**Figure 3.**
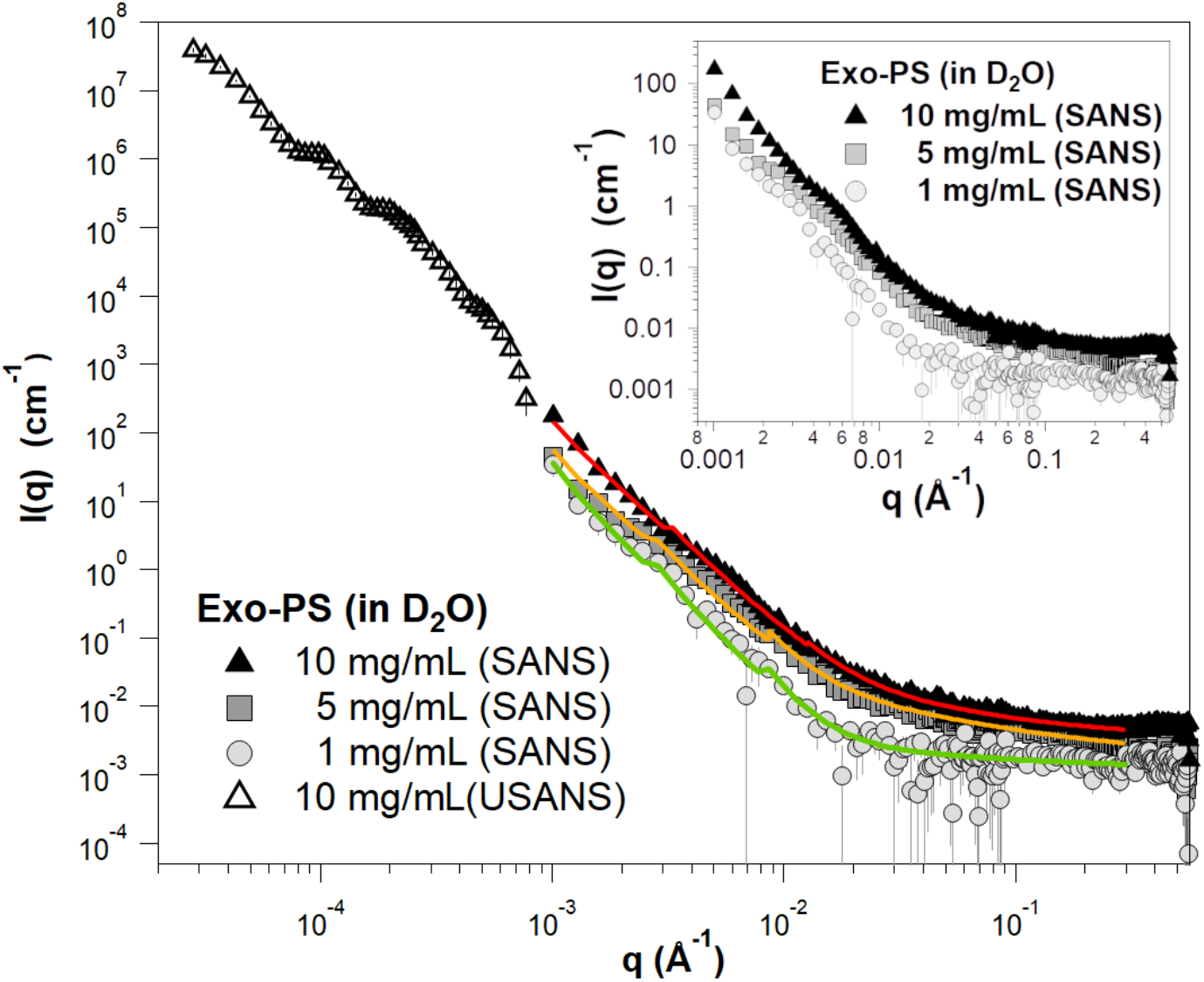
Background-subtracted, reduced SANS, and USANS data for the exo-PS solutions in D2O at varying concentrations. The USANS data shown has been desmeared using the macro- routines for Igor provided by the NCNR to account for the slit-smearing effects of the BT5 instrument on the experimental data. The solid lines depict SANS data fits for the solutions at 1 mg/mL (green), 5 mg/mL (orange), and 10 mg/mL (red) exo-PS; it should be noted that the small discontinuities in the line fit at *q* ≈ 0.03 Å^-1^ and *q* ≈ 0.09 Å^-1^ are a resolution artifact related to the instrumental configurations used for data collection and are not indicative of a sample-related scattering characteristic. The inset provides an enlarged view of the SANS data to highlight the distinctions observed between different concentrations. The error bars represent standard errors derived from counting statistics and, when not visibly discernible, are smaller than the corresponding data markers.

At low *q* values, the SANS data for the 10 mg/mL exo-PS in D_2_O extend down to 0.001 Å^-1^ and exhibit a *q*-dependent intensity profile that reasonably matches the USANS data profile. This indicates the reliability of the desmearing process (a correction applied to the experimental data to account for the slit-smearing effects of the Bonse-Hart USANS instrument) and underscores the consistency between the data collected for the sample in the two scattering regimes^34^.

For *q* values below 0.003 Å^-1^, the SANS scattered intensities of the exo-PS samples at 5 mg/mL and 10 mg/mL display a very similar *q^-n^* dependency characteristic of mass fractals (refer to the Porod exponents in **Table S1**), with *n* ≈ 2.9 reflecting a compact gel structure. An increase in intercluster interactions is observed for the 5 mg/mL and 10 mg/mL exo-PS solutions, as indicated by the larger Porod scales compared to those at 1 mg/mL exo-PS.

The fitting of the exo-PS SANS data produced a consistent correlation length for all concentrations measured (48.9 ± 7.8 Å for 1 mg/mL exo-PS), which is approximately equivalent to the length of four M2 triads (**Figure 4C**). The gelation of exo-PS contributes to the viscosity of the solutions and yields a relatively small correlation length^35^. While the presence of negatively charged glucuronic acid (GlcA) residues can lead to chain-swelling due to electrostatic repulsion, the occurrence of other interactions can stabilize polymer collapse and interchain interactions^23,36,37^. Namely, hydrogen bonds, van der Waals interactions, and ionic bridging which can be promoted by the presence of the divalent cations Mg^2+^ and Ca^2+^ in the culture media^24^. The impact of ionic bridging in polysaccharide structure can be observed by the change in the SAXS profile of exo-PS H2O solutions treated with a chelating agent (see Figure S3).

**Figure 4.**
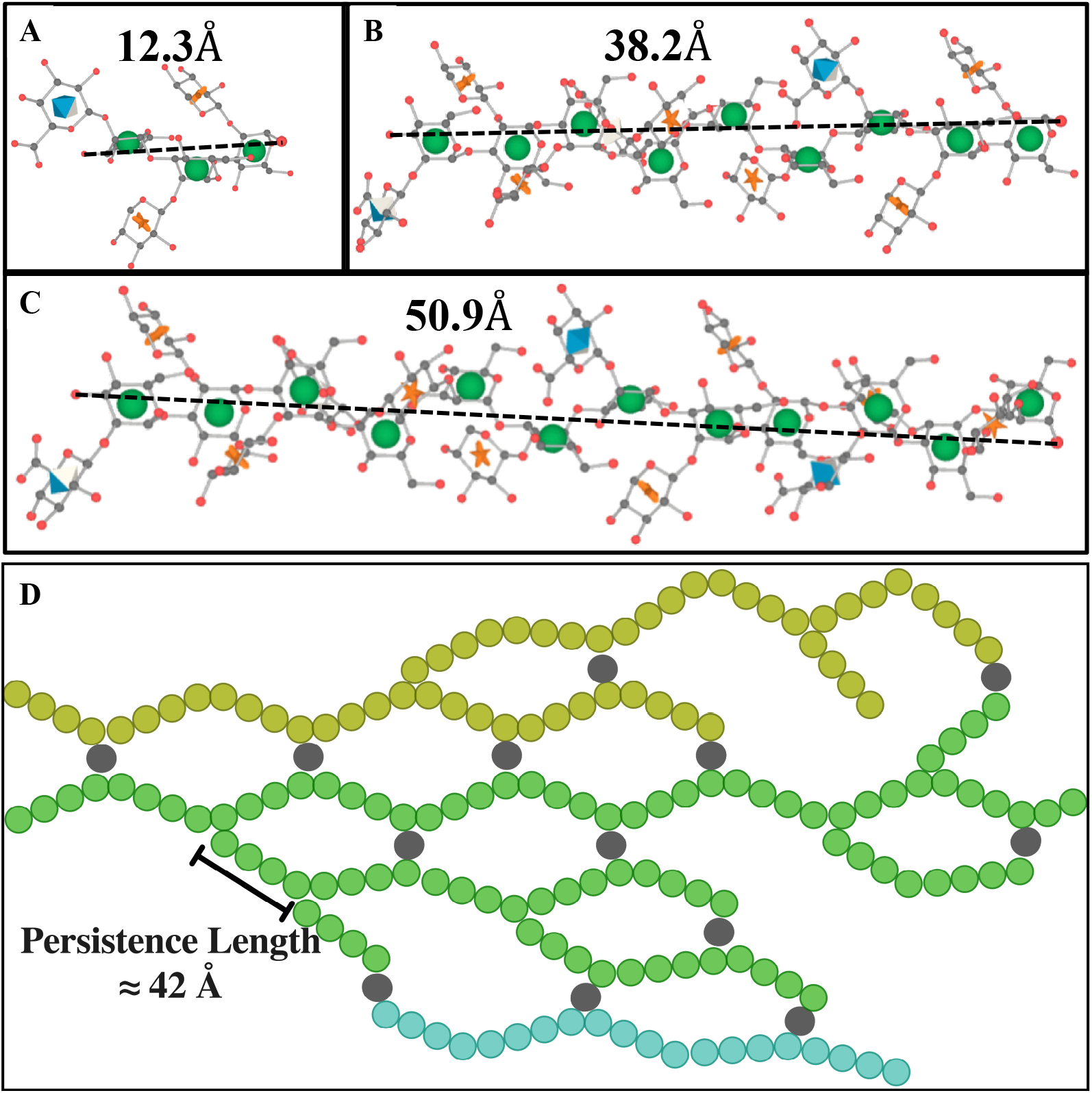
Schematic representation of M2 motif and exo-PS in water for one M2 (**A**), three M2 (**B**), and four M2 (**C**) triads, drawn and energy minimized using GLYCAM^38^. Residues are represented by their symbol nomenclature: green sphere for mannose, orange start for xylose, and blue diamond for GlcA. The distance between the two furthest oxygens that connect mannoses was measured to determine the approximate length of one (12.3 Å), three (38.2 Å), or four (50.9 Å) triads, respectively. (**D**) The exo-PS in water is drawn in 2D to suggest a compatible arrangement of the exo-PS structure, where each circle (except gray circles that represent divalent cations) represents one triad, and three exemplar chains are represented with different colors. A potential pattern of intra- and inter-chain ionic bridging by the divalent cations such as Mg^2+^ and Ca^2+^ (represented by gray circles) present in the cell culture media. The corresponding estimated persistence length (length of the region with rigid rod behavior) is approximately 42 Å. Image created with BioRender.com.

For the 1 mg/mL exo-PS solution, below the overlap concentration, the fit to the SANS data generated a Porod exponent of 3.28 ± 0.16, consistent with a roughness or irregularity of the pores within the gel network, as illustrated by the schematic drawing in Figure 4D and supported by the SEM data at a similar length scale.

For the 5 mg/mL and 10 mg/mL exo-PS solutions, a discernable change in the *q^-n^* dependency of the SANS scattering intensities is observed at *q* values around 0.006 Å^-1^ and beyond, where *n* increases in its value (**Figure 3** inset). In the case of the 1 mg/mL exo-PS solution, however, due to the poorer signal-to-noise ratio, such a transition is not as precisely defined.

The *q*-dependency of the SANS intensity was analyzed at various length scales for the 1 mg/mL exo-PS solution in Figure 5. The Porod region (**Figure 5A**) covers a *q*-range of approximately 0.003 Å^-1^-0.015 Å^-1^, corresponding to structural dimensions within the range of (2π/*q*) ≈ 420 Å- 2090 Å. This is followed by the Lorentz region, characterized by a *q*-dependency of *q*^-1^. The standard Kratky plot (**Figure 5B**) shows that at higher *q* ranges, the profile shifts at *q\** ≈ 0.045 Å^-1^, indicating a rigid rod behavior on a local scale. At this length scale, the PS chain is expected to exhibit rigid rod-like behavior without reorientation or branching^39,40^. The persistence length *l* can be calculated for an ideal Gaussian chain using the formula:

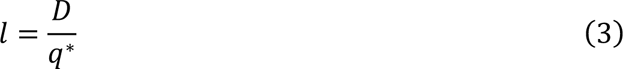

**Figure 5.**
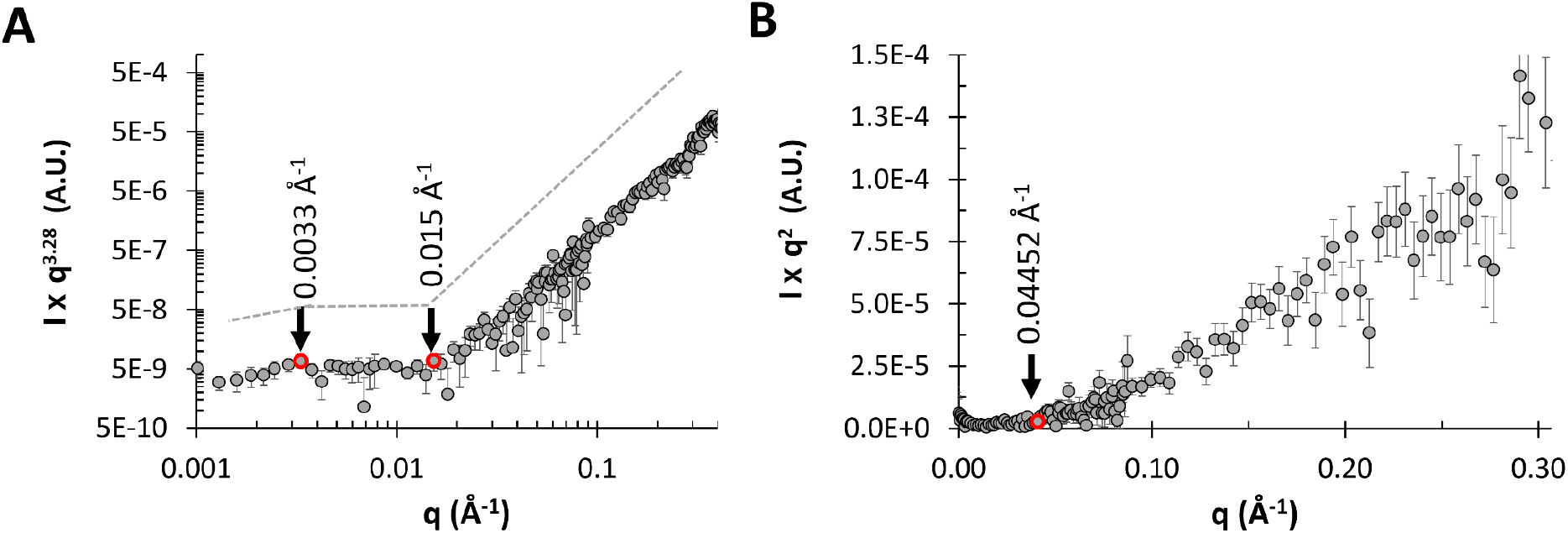
SANS data for 1 mg/mL exo-PS solution in D2O, are shown as (A) Porod exponent- weighted intensities and (B) a standard Kratky plot.

where *D* is a constant with a value of 6/. ≈ 1.91^40^. Applying this approximation to the exo-PS polymer in this local regime, the estimated persistence length is ≈ 42 Å. This length is consistent with that of three or four M2 units, suggesting a rigid rod-like behavior with no interruption between triads within each block.

### Optical Microscopy, SANS, and USANS Analysis of Whole Fungal Cells

Figure 6 and **Table S2** present microscopy of *C. neoformans* H99 cells in different solutions. The microscopic images reveal a relatively high degree of variation in terms of both cell size and capsule thickness for the whole cell and gamma-irradiated cell suspensions. Without gamma irradiation, the average cell diameter was 13.9 ± 2.9 µm in D_2_O and 13.2 ± 3.3 µm in H_2_O, while cells subjected to gamma irradiation had an average cell diameter of 6.4 ± 2.7 µm in D_2_O and 7.2 ± 2.9 µm in H_2_O. The average capsule thickness before gamma-irradiation was also measured: 4.0 ± 1.4 µm in D_2_O and 3.6 ± 1.5 µm in H_2_O. Based on the microscopy data, the choice of solvents did not significantly affect cell di ameter or capsule thickness (unpaired t-test, P > 0.05, ns). However, gamma irradiation removed most of the capsular-PS (unpaired t-test, P < 0.0001, ****).

**Figure 6.**
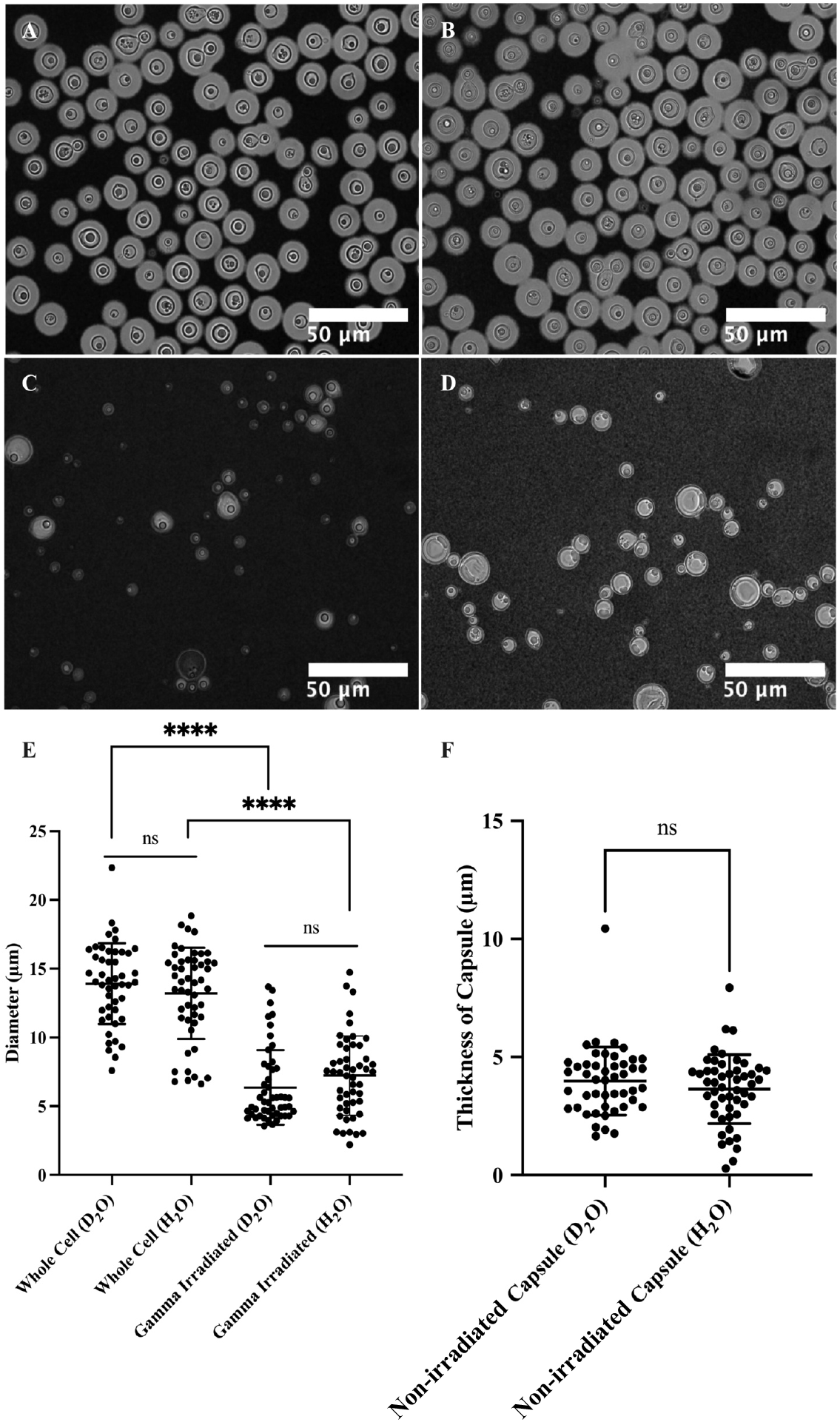
Microscopy of *C. neoformans* H99 cells in water and the effect of gamma irradiation on cellular and capsular dimensions. Samples were counterstained with India Ink particles, which are excluded by the dense PS capsule. Cells without gamma irradiation were resuspended in D2O (**A**) and H2O (**B**), and cells treated with 40 minutes of gamma irradiation were resuspended in D2O (C) and H2O (**D**) as well. The cell diameters of all four samples were estimated (**E**), and capsule thickness for whole cells was obtained as the differences between the radii of cells and cell bodies (G) The t-test analyses are labeled based on their p-value (p > 0.05: ns; p < 0.05: *; p < 0.01: **; p < 0.001: ***; p < 0.0001: ****).

SANS and USANS data on whole and irradiated cells are presented in Figure 7, showing good agreement between the desmeared USANS data and the SANS data. In H2O, the scattering intensities gradually decay with increasing *q* both for irradiated and whole cell samples, reaching similar incoherent scattering background intensities of approximately 0.07 Å^-1^-0.08 Å^-1^, dominated by contributions to incoherent scattering from the hydrogen atoms present. In contrast, D2O samples exhibit a substantially lower incoherent neutron scattering background, allowing a discernable difference in the SANS profiles to emerge at *q* > 0.1 Å^-1^. This range corresponds to the expected contribution of the M2 triads towards scattering. Notably, at these higher *q* values, gamma-irradiated samples display a distinct profile with a decrease in scattering intensities. This observation is consistent with the disruption of the M2 triads that SANS detects for whole cells in D_2_O, but not in H_2_O where it was observed that the exo-PS scattering length density is matched out. Differences are also evident in the Kratky plots for samples measured in D_2_O, as present in Figure S5 in the supporting information.

**Figure 7.**
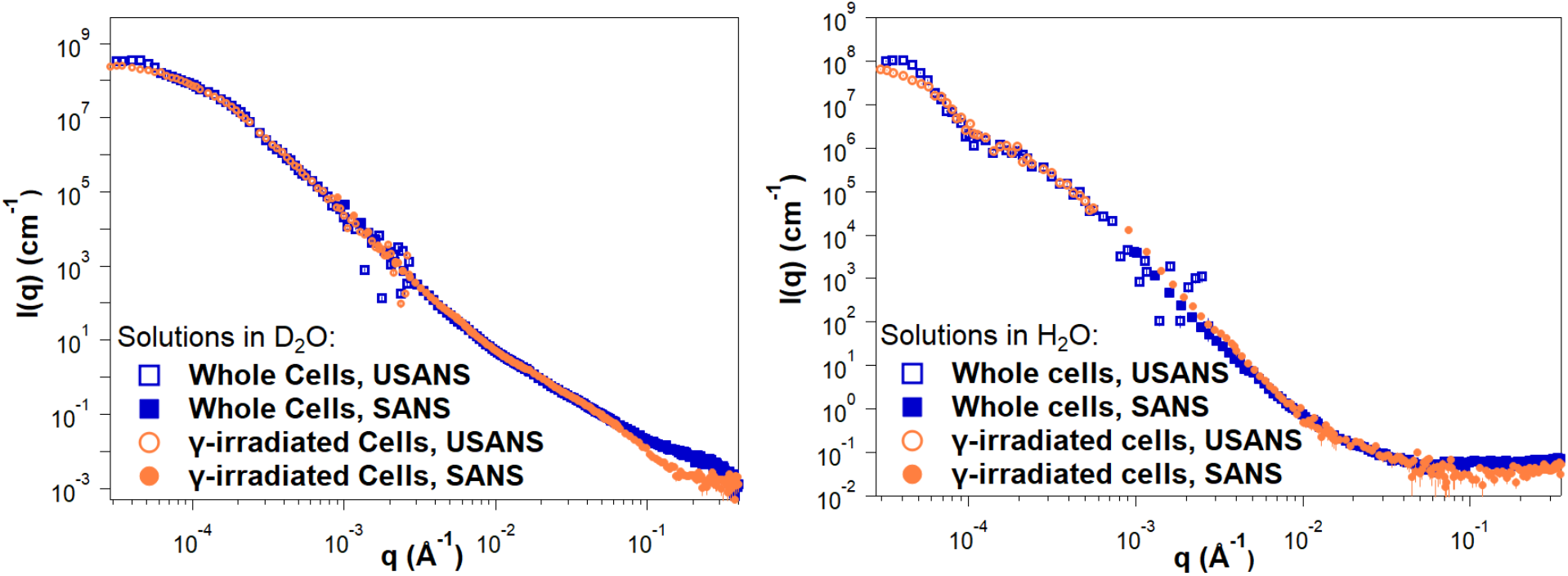
Reduced SANS (buffer subtracted) and USANS data collected for fungal cells in D2O and H2O solutions, for both intact and gamma-irradiated cells. The desmeared USANS data displayed compensates for the slit-smearing effects on the experimental data, allowing direct comparison with the SANS data for each sample. Error bars represent standard errors from counting statistics and are smaller than the corresponding data marker when not visible.

Given the high polydispersity of the samples, USANS data fitting was not attempted, as neutron scattering probes a significant amount of bulk sample compared to the images in Figure 6. Additionally, unknown contributions to the scattering profile from cellular components such as the nucleus or the cell wall further complicate data fitting. The difference in size between whole and gamma-irradiated cells is apparent from the USANS profiles for both H_2_O and D_2_O samples, where the scattering profiles show a maximum intensity plateau at *q* < 0.00005 Å^-1^, as the intensities reach the Guinier regime. The calculated radii of gyration for irradiated cell samples, derived from Guinier analysis (**Figure S4** and **Table S3**), are consistent with the dimensions observed by optical microscopy. For non-irradiated samples, insufficient data points were collected in the Guinier regime, preventing an unambiguous calculation of the corresponding larger cell radii of gyration.

Since exo-PS in H_2_O solutions did not exhibit measurable contrast in the SANS regime, even at concentrations of 10 mg/mL, the discernible discrepancies in cell size between whole and gamma-irradiated cells in H_2_O indicate differentiation between the exo-PS and capsular-PS. This is consistent with a gradient of PS densities in the pristine whole cells, ranging from an outer, more hydrated layer to an inner, denser layer closer to the cell wall, which is less solvent accessible and more resistant to ablation^23^.

In the USANS data for H_2_O solutions, there is an inflection point at *q* ≈ 0.0001 Å^-1^ that is absent in the samples containing D_2_O. Considering that no significant increase in polydispersity was detected by optical microscopy for the samples in D_2_O compared to H_2_O, the absence of inflection in D_2_O solutions is not consistent with a resolution effect. Instead, the data may reflect a structural characteristic of the fungal cell body for which the solutions in H_2_O provide better SLD contrast. Given the complexity of the fungal cell and the unknown SLD of the different cell components, no specific structure or organelle can be objectively assigned to this area of the USANS profile.

### Fractal Analysis of Scanning Electron Micrographs

Image analysis of encapsulated whole cells and capsular structures reveals fractal patterns with dimensions ranging from 1.6 to 1.85 (Figure 8). Despite these samples undergoing dehydration during SEM processing and thus not being in their native state, the presence of fractal patterns in the polysaccharides is consistent with the neutron scattering data in solution. Moreover, this fractal pattern was observed even after capsule dehydration and the coalescing of PS molecules into thick fibrils. This suggests a connection between the hydrated and dehydrated structures of polysaccharides, possibly reflecting a residual capsular resistance to dehydrating conditions.

**Figure 8.**
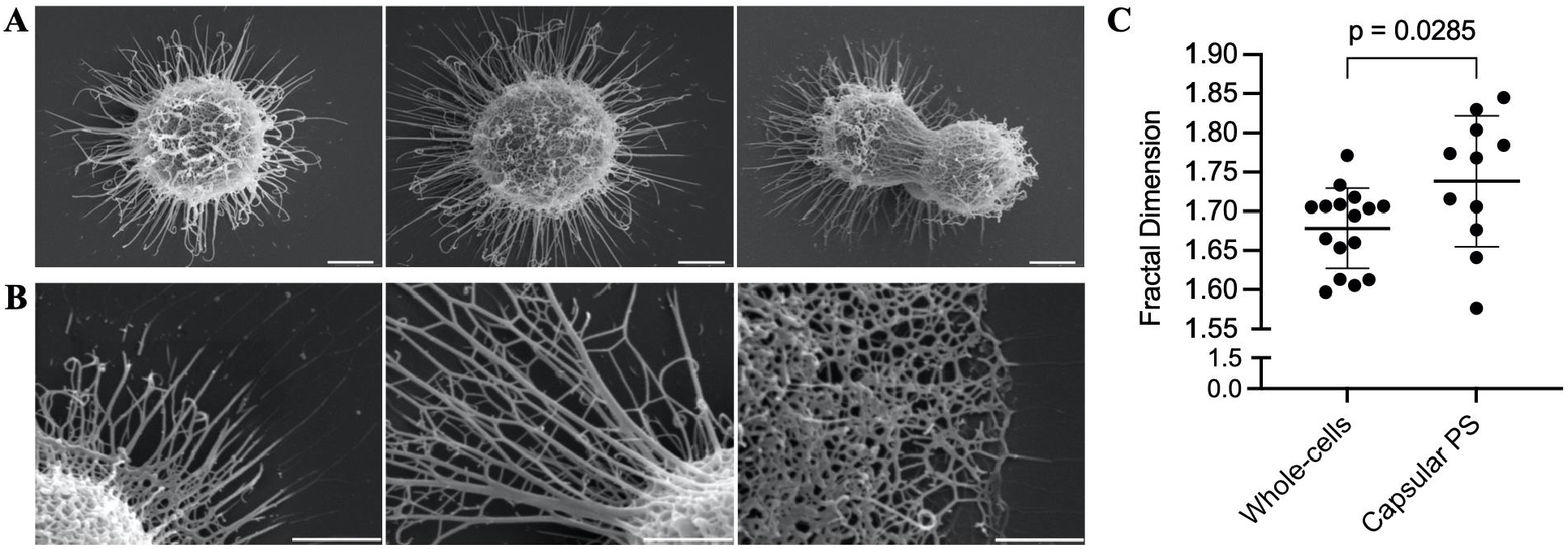
Fractal analysis of SEM micrographs of whole encapsulated cells and capsular-PS structures. (A) Representative whole encapsulated *C. neoformans* cells and (B) capsular-PS structures, showing the fractal structure and the presence of an irregular pore network. Scale bars represent 2 and 1 micrometers, respectively. (C) Fractal dimensions of 27 micrographs were analyzed using the FracLac plugin in ImageJ.

## CONCLUSIONS

This study demonstrates the effective use of neutron scattering and different neutron scattering contrasts (utilizing solutions with 0% and 100% D_2_O) to gain insights into the structural characteristics of both *C. neoformans* exo-PS and cells under their native conditions. Our findings present compelling evidence that exo-PS inherently exhibits mass fractal characteristics, representing a self-similar branched system or network spanning a wide range of size scales. While the presence of fractals in certain polysaccharides is not uncommon, the specific characteristics of these fractals, such as fractal dimension, can vary widely due to multiple factors, such as molecular composition, surrounding environment, and processing conditions^6,22^.

The SANS results from exo-PS scattering are consistent with a collapsed chain-like behavior stabilized by interchain and intrachain interactions, including divalent cation bridges between negatively charged glucuronic acid residues, a phenomenon previously reported in the presence of water molecules^25^. Future SANS and USANS studies should address the concentration effects of exo-PS on hydrogen-to-deuterium exchange levels and buffer accessibility (including chelating agents if used to further investigate the role of cation bridging in the structuring of the polysaccharide). High concentrations are likely to impact solvent buffer accessibility to the polysaccharide, while lower concentrations may favor more uniformly chain-hydrated states and minimize scattering effects arising from overlapping and entanglement.

The observed rigid rod behavior of exo-PS at local scales, with an estimated persistence length of approximately 42 Å, suggests that the arrangement of three to four GXM triads, particularly the M2 motif, involves short repeats where the chain direction may change at the end of each repeat, resulting in an overall semi-flexible structure. This local-scale rigidity of exo-PS agrees with molecular modeling studies, which propose that, at least within six GXM motifs, the ends of the chain do not bend into close proximity^41^.

As a dominant virulence factor often targeted for antibody treatment, exo-PS plays a crucial role in infection. Recent research has identified decasaccharide (serotype A) as a possible minimal size for effective neutralizing mAb recognition, but our data offer a broader range of oligosaccharide sizes suitable for testing immune responses^42^. To better characterize the exo-PS secreted in humans during infection, similar studies on exo-PS secreted by isolated infecting fungal cells cultured in media with the cation compositions of human fluids should be conducted to explore the effects of different types and concentrations of counterions.

It is important to note that sample preparation protocols can influence the measured structural properties, as previously suggested^10^. The experimental data reflects only the structural features of the selected sample and it is likely that the parameters measured here would vary with different types of preparations. Varying molar mass cutoffs during filtration can result in significant differences in the USANS regime (data not shown) and previous studies have shown that *C. neoformans* GXM fractions of different molar masses are functionally distinct^43^. Further work is needed to systematically characterize USANS and microscopy data for exo-PS chains representing the range of molar masses found *in vivo*. Lastly, future SANS studies could utilize isotope-labeled dextrose or specific precursors to enhance the contributions of capsular-PS to the overall scattering profiles measured from whole cells, with the goal of further elucidating the differences between capsular-PS and exo-PS^10^.

In summary, the application of neutron scattering to *C. neoformans* polysaccharide provides important new insights into its structure, including evidence for a fractal-like structure arising from intermolecular interactions of semi-rigid PS segments. This arrangement forms a mesh that emerges as the capsular structure observable in the macromolecular world. This in turn raises the exciting possibility that the capsule emerges from perhaps a few local interactions between polysaccharide molecules, guided by specific local rules, resulting in the magnificence that is the visible *cryptococcal* capsule. The capsule is a highly hydrated structure that is difficult to study directly using available techniques. Therefore, our understanding of the capsule is derived from synthesizing information gathered from techniques at differing scales, spanning from microscopic observations to neutron scattering and the chemistry of the individual sugars, in the creation of testable models. In this regard, the addition of neutron scattering to the methodologies used for studying *cryptococcal* polysaccharide introduces a new and welcomed analytical tool.

## ASSOCIATED CONTENT

**Supporting Information**: dynamic light scattering analysis of exo-PS, neutron, and X-ray scattering data and data fitting for exo-PS samples, microscopy data for whole cell samples, and Guinier fittings and Kratky plots for the fungal cell samples based on USANS and SANS data, respectively (PDF).

## AUTHOR INFORMATION

### Author Contributions

RJBC and SCMT designed the neutron scattering experimental approach. CS and RJBC designed the cell culture and polysaccharide isolation protocols, prepared samples, and collected optical microscopy data. SCMT and CS collected and reduced the neutron scattering data. SCMT and ZW carried out the data fitting and analyses. AB carried out the fractal dimension analysis on SEM images. The manuscript was written through the contributions of all authors. All authors have approved the final version of the manuscript.

### Notes

The authors declare no competing financial interest.

## ACKNOWLEDGMENT

This work benefited from the use of the SasView application, originally developed under NSF award DMR-0520547. SasView contains code developed with funding from the European Union’s Horizon 2020 research and innovation program under the SINE2020 project, grant agreement No 654000. SCMT is grateful for funding from the cooperative agreement #70NANB20H133 from NIST, U.S. Department of Commerce. We acknowledge the support of the National Institute of Standards and Technology, U.S. Department of Commerce, in providing the neutron research facilities used in this work. This work utilized facilities supported in part by the National Science Foundation under Agreement No. DMR-0944772. Certain commercial equipment, software, instruments, and materials are identified to foster understanding. Such identification does not imply recommendation or endorsement by the National Institute of Standards and Technology, nor does it imply that the materials or equipment identified are necessarily the best available for the purpose. The statements, findings, conclusions, and recommendations are those of the authors and do not necessarily reflect the view of NIST or the U.S. Department of Commerce. RJBC was supported by the Johns Hopkins University Center for AIDS Research (P30AI094189). The authors also thank Dr. Scott A. McConnell for reviewing the data and providing valuable input.

## ABBREVIATIONS

PS, polysaccharide; GXM, Glucuronoxylomannans; SANS, small-angle neutron scattering; USANS, ultra-small-angle neutron scattering; SEM, scanning electron microscopy; SLD, scattering length density.

