## Supporting Information for "Neutron Scattering Analysis of *Cryptococcus neoformans* Polysaccharide Reveals Solution Rigidity and Repeating Fractal-like Structural Patterns"

Ziwei Wang[1], Susana C. M. Teixeira[2,3*], Camilla Strother[1], Antony Bowen[1], Arturo Casadevall[1], and Radamés JB Cordero[1*]

[1]Molecular Microbiology and Immunology, Johns Hopkins Bloomberg School of Public Health, Baltimore, Maryland, 21205, USA.
[2]NIST Center of Neutron Research, National Institute of Standards and Technology, Gaithersburg, Maryland, 20899, USA.
[3]Department of Chemical and Biomolecular Engineering, University of Delaware, Newark, Delaware, 19716, USA.
[*]Corresponding authors.

### 1- Dynamic Light Scattering Data on the Exo-PS Sample.

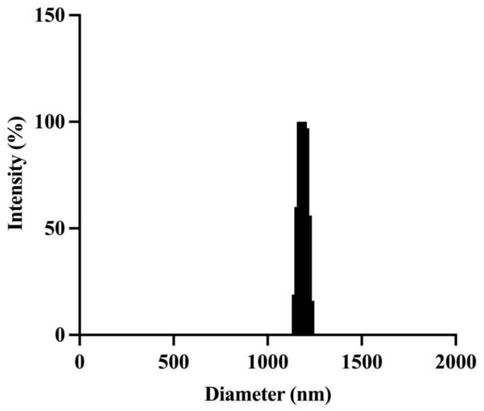


**Figure S1.** Dynamic light scattering (DLS) analysis of 10 mg/mL exo-PS solution accumulated on a 10 kDa molar mass cutoff membrane. The polydispersity of the exo-PS was 0.355, determined by DLS coupled with a 90Plus/BI-MAS Multi-Angle Particle Sizing analyzer (Brookhaven Instruments Corp., NY, USA).

### 2- Neutron Scattering Data on the Exo-PS Samples and Data Fitting.

**Figure S2.** Background subtracted SANS data – data collected for buffer-only solutions was subtracted from the PS-containing samples – measured on exo-PS in $H_2O$ at 30 °C and various concentrations, as labeled. The scattered intensities show approximately flat profiles, indicative of poor contrast for the scattering angles measured. The error bars represent standard errors from counting statistics and when not visible are smaller than the corresponding data marker.

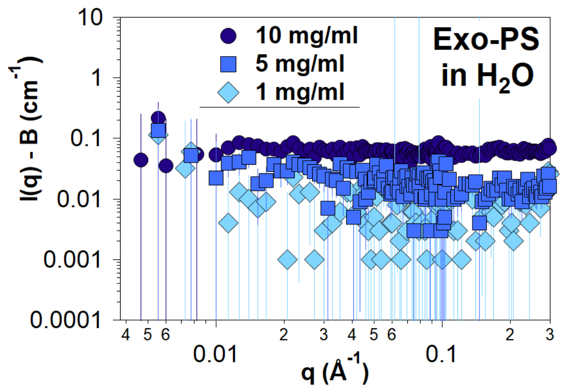

Small angle neutron scattering (SANS) data on the exo-PS solutions in $D_2O$ were fitted using the correlation length model available in SasView[1], as described in the main document, where the calculated scattered intensities are defined as[2]:

$$I(q) = \frac{A}{q^n} + \frac{C}{1 + (q\xi)^m} + B$$

where the first term describes Porod scattering from pore clusters (exponent $n$) and the second term is a Lorentzian function describing scattering from polymer chains (exponent $m$). The second term characterizes the polymer/solvent interactions, and the two multiplicative factors $A$ and $C$ are, respectively, the Porod scale and the Lorentz scale. $\xi$ is a correlation length for the polymer chains, and $B$ is a $q$-independent incoherent neutron scattering intensity contribution to I($q$). The calculated intensities from the correlation length model were smeared to match the instrumental pinhole smearing read from the reduced experimental data file. Fits to the SANS data minimize a statistical parameter that quantifies the differences between observed (measured) data and theoretical (calculated from the correlation length model) data points, defined as:

$$\chi^2 = \sum \frac{(Y_{obs} - Y_{theor})^2}{weight^2}$$

where the weight given to each data point takes into account the counting errors associated with the measured intensities, read from the data file. The goodness-of-fit was assessed using a reduced chi-squared parameter that takes into account the degrees of freedom (the number of observed points $N_{obs}$ subtracted by the number of fitted parameters $N_{fitted}$):

$$\chi^2_{red} = \frac{\chi^2}{N_{obs} - N_{fitted}}$$

**Table S1.** Fitting parameters for the exo-PS SANS data using a modified correlation length model[2]. The data were fitted for $0.001 \leq q$ (Å$^{-1}$) $\leq 0.3$. Higher $q$ data were not included to prevent biasing the fits with data points that are sensitive to background subtraction effects during data reduction. Values where no uncertainties are shown were kept fixed during the fits.

| **Exo-PS ($D_2O$)** | **1 mg/mL** | **5 mg/mL** | **10 mg/mL** |
|---|---|---|---|
| ***q*-range (Å$^{-1}$)** | 0.001 – 0.30 | 0.001 – 0.30 | 0.001 – 0.30 |
| **Reduced Chi-squared** | 0.83 | 1.74 | 1.00 |
| **Volume fraction** | 0.001 | 0.005 | 0.01 |
| **Background *B* (cm$^{-1}$)** | 0.0012 | 0.0019 | 0.0035 |
| **Lorentz scale *C*** | 2.2 ± 0.5 | 3.0 ± 0.1 | 2.0 ± 0.1 |
| **Porod scale *A*** | $(3.2\pm1.5)\times10^{-6}$ | $(16\pm0.3)\times10^{-6}$ | $(18.0\pm2.5)\times10^{-6}$ |
| **Correlation length $\xi$ (Å)** | 48.9 ± 7.8 | 48.6 ± 1.1 | 49.9 ± 1.3 |
| **Porod exponent *n*** | 3.28 ± 0.16 | 2.89 ± 0.04 | 2.91 ± 0.02 |
| **Lorentz exponent *m*** | 0.79 ± 0.16 | 1.02 ± 0.03 | 1.06 ± 0.04 |

### 3- Small Angle X-ray Scattering Data on Exo-PS Samples.

Preliminary Small Angle X-ray Scattering (SAXS) experiments were conducted on samples dialyzed against a buffer containing a chelating agent (10 mM NaCl and 1 mM EDTA in $H_2O$). The data (see **Figure S3**) was compared for samples prepared in pure water, and samples treated by sonication for 20 minutes prior to data collection (a common protocol used to disaggregate polysaccharides, see for example presented by Geresh *et al.*[3]).

There is insufficient data at low $q$ to accurately measure the $q$-dependency of the intensities and the scale of the fractal behavior but, compared to exo-PS prepared in water only, the results show a clear shift in the scattering intensities for the EDTA-treated sample, particularly at $q \approx 0.025$ Å$^{-1}$ and higher, where the data is consistent with significant disaggregation of the polysaccharide (comparable to the effects of sonication).

While this preliminary data supports the theory of a strong role for cationic bridging in the structuring of exo-PS, this comparison should take into consideration a few limitations, namely: (1) potentially restricted accessibility of EDTA and solvent to parts of the compact gel network and (2) the known susceptibility of exo-PS to ionizing radiation-induced ablations (that can occur from exposure to X-rays during SAXS measurements), particularly if prior sonication increases free-radical diffusion and accessibility of the polymer[4].

A deeper understanding of the role of cation bridging will require a series of systematic studies using optimized sample treatment protocols to assess and mitigate radiation damage (SAXS), and neutron scattering data (SANS) to prevent potential sample discrepancies.

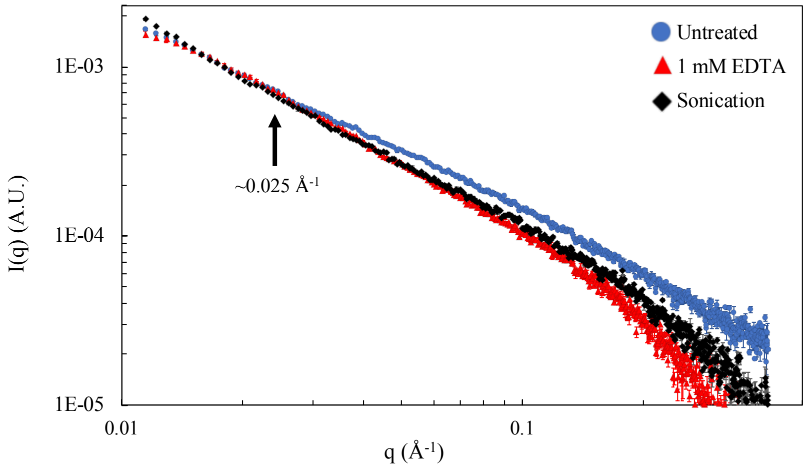


**Figure S3.** SAXS data collected of 10 mg/mL exo-PS samples prepared in pure water, in a buffer containing EDTA as a chelating agent (EDTA treated) and for samples that were sonicated for 20 minutes prior to data collection (Sonicated). Buffer contributions have not been subtracted from the shown intensity profiles. The data points are colored as described in the legend. Error bars represent standard errors from counting statistics. Data were collected at the Brazilian Synchrotron Light Laboratory (LNLS) with SAPUCAIA beamline (Scattering APparatUs for Complex Applications and *In-situ* Assays).

### 4- Light Microscopy, USANS, and SANS Data on Whole Fungal Cells

**Table S2** shows estimates of the cell dimensions based on optical microscopy data, as well as estimates of the corresponding radii of gyration $R_g$, assuming an approximately spherical shape with a uniform distribution of neutron scattering length density. The radius of gyration for a sphere can be estimated from the real-space radius $R$ of the cells using the formula:

$$R_g^2 = \frac{3}{5}R^2$$

**Table S2.** Cell dimensions are estimated from optical microscopy data of samples containing whole and gamma-irradiated cells in different buffers, where $D$ is the diameter, $V$ is the volume, $R_g$ is the expected radius of gyration, and $T$ is the thickness.

| **Cell Samples** | | **Cell** | | | **Cell Body** | | **PS Capsule** | |
|---|---|---|---|---|---|---|---|---|
| | | **D (µm)** | **V ($\mu m^3$)** | **$R_g$ (µm)** | **D (µm)** | **V ($\mu m^3$)** | **T (µm)** | **V ($\mu m^3$)** |
| **Whole** | $D_2O$ | $13.9 \pm 2.9$ | 1406 | $5.4 \pm 1.9$ | $5.9 \pm 1.8$ | 109 | $4.0 \pm 1.4$ | 1297 |
| | $H_2O$ | $13.2 \pm 3.3$ | 1205 | $5.0 \pm 1.6$ | $5.9 \pm 1.5$ | 109 | $3.6 \pm 1.5$ | 1096 |
| **Irradiated** | $D_2O$ | $6.3 \pm 2.7$ | 135 | $2.3 \pm 1.4$ | $6.4 \pm 2.7$ | 134 | - | - |
| | $H_2O$ | $7.2 \pm 2.9$ | 188 | $2.7 \pm 1.5$ | $7.2 \pm 2.9$ | 188 | - | - |

For both whole and gamma-irradiated cells, the USANS intensities appear to trend towards a plateau value at low $q$, reflecting the size of the particles in solution. Assuming that there are no significant interactions between the cells at the concentrations used and that the cells are approximately spherical with a uniform distribution of neutron scattering length density, the Guinier approximation is valid for $q \times R_g < 1$:

$$ln\ I(q)\ = ln\ I(0)\ - \frac{q^2 Rg^2}{3}$$

where I($q$) are the scattered intensities as a function of the momentum transfer magnitude $q$[5,6]. Guinier fits for the irradiated cell samples are shown in **Figure S4**. As shown in **Table S3**, although the USANS data captured part of the Guinier regime, given that USANS data were collected for $q > 3 \times 10^{-5}$ Å$^{-1}$, there are insufficient data points in the Guinier regime for the non-irradiated cells.

**Table S3.** Parameter fits for the USANS data on the gamma-irradiated cells, where the slit-smearing effects of the experimental resolution were taken into account when performing the Guinier fits, and *I(0)* is the intensity at zero $q$. The errors shown report the standard deviation of the parameters (square root of the covariance matrix diagonal elements), and $r^2$ is the linear regression goodness-of-fit.

| **Cell samples** | **$R_g$ (µm)** | **I(0) $\times 10^8$ (A.U.)** | **$q \times R_g$** | ***q*-range $\times 10^{-5}$ (Å$^{-1}$)** | **$r^2$** |
|---|---|---|---|---|---|
| Whole, $D_2O$ | - | - | < 1 | < 1.9 | - |
| Whole, $H_2O$ | - | - | < 1 | < 2 | - |
| Irradiated, $D_2O$ | $2.063 \pm 0.031$ | $(2.76 \pm 0.02)$ | 0.59 - 1.28 | 3 - 6 | 0.956 |
| Irradiated, $H_2O$ | $3.168 \pm 0.498$ | $(0.85 \pm 0.09)$ | 0.85 – 1.12 | 3 - 4 | 0.502 |

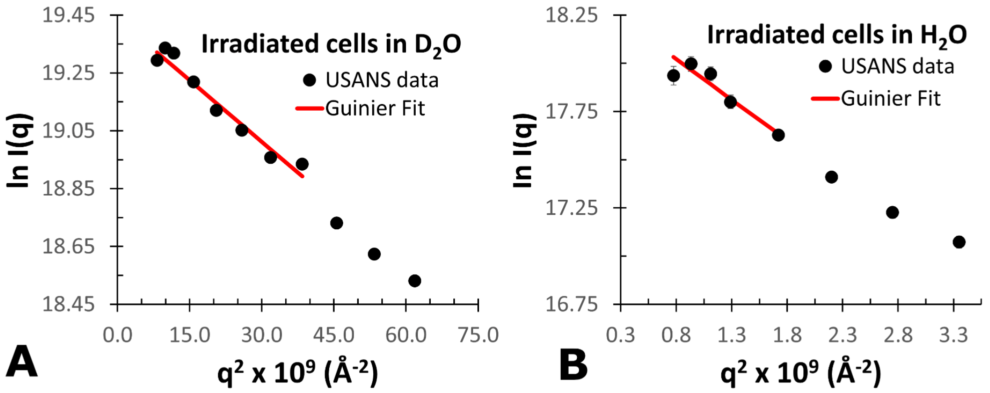


**Figure S4.** Guinier plots and corresponding data fits for the irradiated cell samples in **(A)** $D_2O$ and **(B)** $H_2O$ buffers. Error bars represent standard errors from counting statistics and when not visible are smaller than the corresponding data marker.

In the SANS regime, contributions to the scattering from multiple components of the fungal cells preclude a more objective analysis of the data in terms of the structure of the capsular PS. At high-*q*, however, the differences observed between whole and irradiated samples in the presence of $D_2O$ are consistent with the branched structure of the polysaccharide contributing to scattering from the whole cells, and absent in the irradiated samples. The differences at high-*q* for whole and irradiated samples are not observed in $H_2O$, where exo-PS data had already shown that the PS is matched out.

A Kratky plot calculated from the scattering data for the solutions in $D_2O$ is shown in **Figure S5**, where the differences between whole and gamma-irradiated cell samples are highlighted. The Kratky plot does not show the rigid rod-like profile for the irradiated samples. Although it should be highlighted that Kratky plots are extremely sensitive to effects of inconsistencies in background subtraction, and potential contributions from other materials present in the fungal cells cannot be ruled out, the data is consistent with the removal of capsular PS from the irradiated cells, as observed in the light microscopy images.

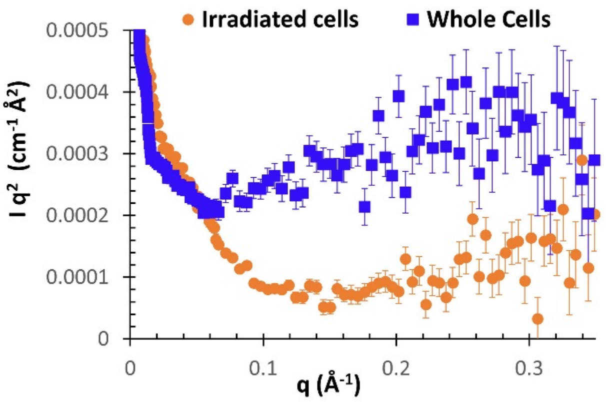


**Figure S5.** Kratky plots calculated from the SANS data collected on fungal cell samples in $D_2O$. Error bars represent standard errors from counting statistics and when not visible are smaller than the corresponding data marker.
